# Impact of the elderly lung mucosa on *Mycobacterium tuberculosis* metabolic adaptation during infection of alveolar epithelial cells

**DOI:** 10.1101/2024.05.06.592753

**Authors:** Angélica M. Olmo-Fontánez, Anna Allué-Guardia, Andreu Garcia-Vilanova, Jeremy Glenn, Shu-Hua Wang, Robert E. Merritt, Larry S. Schlesinger, Joanne Turner, Yufeng Wang, Jordi B. Torrelles

## Abstract

**Background:** Tuberculosis (TB) is one of the top leading causes of death due to a single infectious agent. Upon infection, *Mycobacterium tuberculosis* (*M.tb*) is deposited in the alveoli and encounters the lung mucosa or alveolar lining fluid (ALF). We previously determined that increased *M.tb* replication in human macrophages and alveolar epithelial cells (ATs) is mediated by age-associated changes in human ALF. Here we determine the transcriptional profile of *M.tb* when exposed to healthy ALF from adult (A-ALF) or elderly (E-ALF) individuals before and during infection of ATs.

**Results:** Prior to infection, exposure to E-ALF upregulates *M.tb* genes associated with the ESX-4 secretion system, immunomodulatory proteins from the ESX-5 system, and genes encoding phospholipases, phosphatases, and proteases. During infection, E-ALF exposure upregulates key *M.tb* genes from the ESX-5 secretion system, genes associated with PDIMs biosynthesis and transport, and genes linked to bacterial oxidative stress defense mechanisms.

**Conclusions:** These findings demonstrate how altered ALF in old age can impact the metabolic status of *M.tb*, enabling greater adaptation to the host and potentially explaining *Mtb’*s increased survival within host cells. Importantly, we present the first transcriptomic analysis on the impact of the elderly lung mucosa on *M.tb* pathogenesis during intracellular replication in ATs.

## BACKGROUND

The increase in worldwide elderly population (60 years and older) is projected to double to 2.1 billion by 2050 (1), and close to 95 million by 2060 in the United States alone (2). Unfortunately, living longer is associated with ongoing molecular and physiological changes of natural lung aging, leading to decreased lung function and weakened immune responses, increasing susceptibility to lung diseases and respiratory infections (3–7). Consequently, the elderly population is at high risk of contracting respiratory diseases, including tuberculosis (TB) (5, 8).

More than 4,300 people die from TB daily, making it a top infectious killer globally (9). TB is caused by airborne pathogen *Mycobacterium tuberculosis* (*M.tb*), spread mainly through inhalation and deposited into the distal portion of the airways and alveoli. In this environment, *M.tb* comes in contact with the lung mucosa or alveolar lining fluid (ALF), which contains soluble innate factors such as surfactant proteins A and D (SP-A/SP-D), complement, hydrolytic enzymes, antimicrobial peptides, and others, which can trigger subsequent innate and adaptive immune responses (10, 11). The increased susceptibility of the elderly to TB may be attributed to age-related alterations in the lung environment, including innate soluble components of the ALF. Our research has shown that aging impacts the pulmonary environment by causing a shift towards a pro-inflammatory and pro-oxidative state, thus reducing the functionality of soluble innate immune proteins such as SP-A and –D (12–14). Age-associated alterations in ALF components and their functional status are linked to decreased ability of phagocytes to control *M.tb* infection *in vitro* and *in vivo* (13, 15).

Alveolar epithelial type cells (ATs) are non-professional phagocytes that line the alveolar epithelium forming a physical barrier, preventing microbial invasion and directly contributing to host defense against *M.tb* infection (11, 16, 17). ATs consist of two distinct cell types; type I (ATIs), the most prevalent cell type that gives the alveolus its structured shape and permits gas exchange (16), and type II (ATIIs), which play an essential role in maintaining alveolar homeostasis (11, 18). Our previous studies showed that *M.tb* exposure to elderly ALF promotes increased *M.tb* replication and growth within ATs, linked to altered intracellular trafficking and resulting in bacterial translocation into the ATs cytosol (15), dampening host protective immune responses.

Here, we first aimed to define *M.tb* metabolic adaptations upon exposure to different human ALF environments [adult (A)-ALF: 18-45 years of age; and elderly (E)-ALF: >69 years of age] prior to infection. Then, we determined the transcriptional status of A- and E-ALF-exposed *M.tb* during infection in ATs. We and others have shown *M.tb*’s ability to escape into the host cytosol (15, 19–23), hence we particularly focused on genes related to the ESX system and PDIM production as potential strategies of bacterial translocation, as well as other genes associated with *M.tb* pathogenicity. Our findings indicate that exposure of *M.tb* to E-ALF alters its transcriptional profile prior to infection, but also enhances the expression of virulence-associated genes (mainly the ESX secretion system and oxidative stress defense mechanisms) during infection, allowing *M.tb* to better adapt for survival within host cells.

## RESULTS

### Differential expression analysis of ALF-exposed *M.tb* prior to infection

Our previous studies demonstrated that when *M.tb* contacts the human lung mucosa, ALF hydrolytic enzymes modify the *M.tb* cell envelope, eliciting innate and adaptive immune responses that impact subsequent *M.tb* infection outcomes *in vitro* and *in vivo* (10, 13, 24). We recently reported that exposure to E-ALF drives *M.tb*’s translocation into the cytosol, favoring its replication and growth in ATs (15). Based on these findings, we explored the transcriptional changes that *M.tb* experiences after being exposed to human A-ALF or E-ALF at their physiological concentration within the lung. We focused on the following mechanisms reported to play a role in *M.tb* escape into the cytosol: i) ESX secretion system, ii) Phospholipases, iii) PDIMs biosynthesis, iv) Others, including pathways reported to play a role in cytosol translocation of other pathogens. Experimental conditions were designated as A-ALF-exposed *M.tb* (A, pool of n=4 different human donors), E-ALF-exposed *M.tb* (E, pool of n=4 different human donors) and unexposed *M.tb* (UE) (**Fig. 1A**). To determine the effects of ALF exposure on *M.tb* prior to infection, we performed differential expression (DE) analysis (log_2_ FC equal or greater than an absolute value of 1 and FDR < 0.1) of ALF-exposed *M.tb* (A-ALF or E-ALF) compared to unexposed *M.tb* (A *vs*. UE and E *vs*. UE comparisons, respectively, **Fig. 1A**).

**Figure 1.**
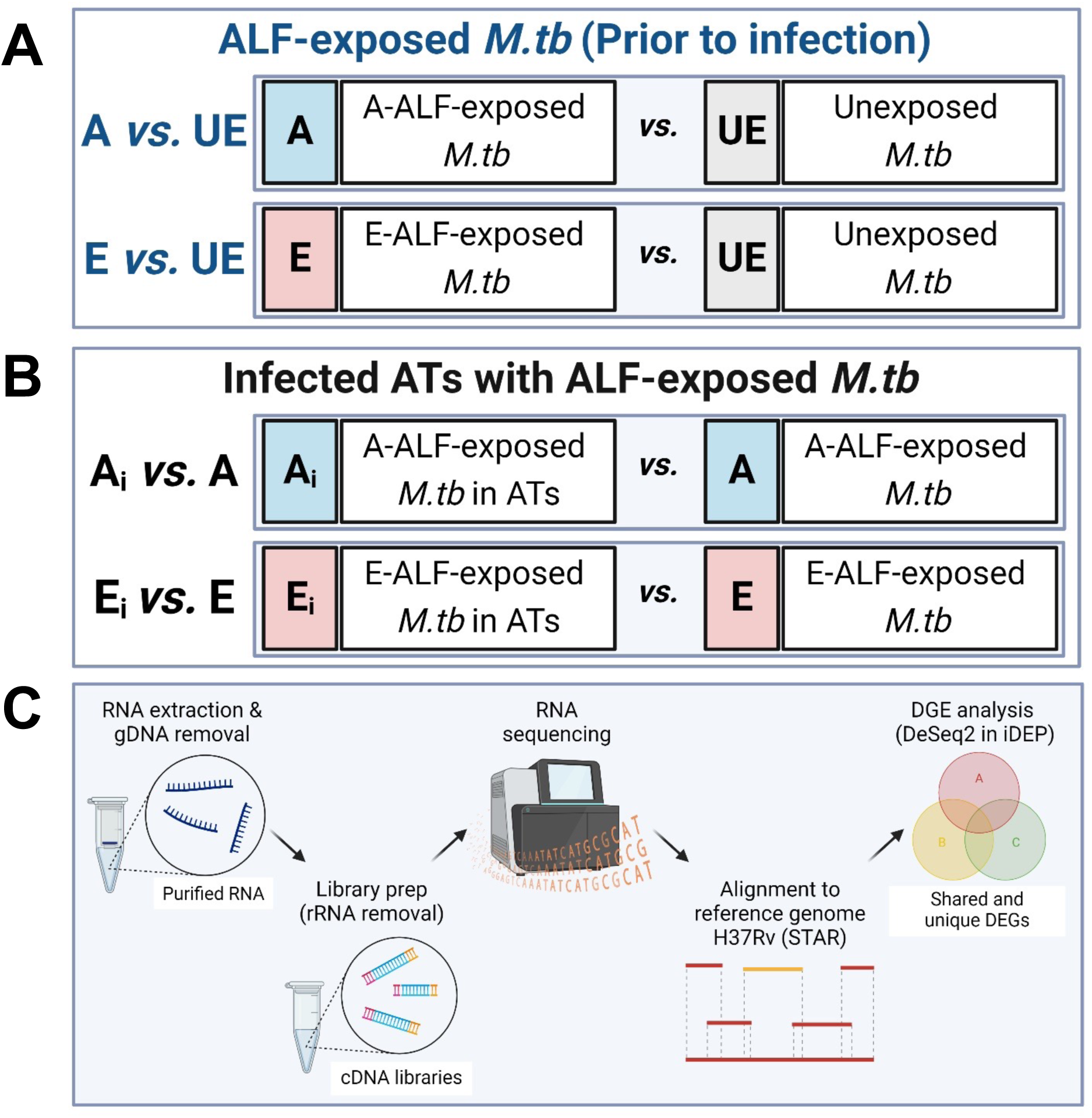
Illustration of the experimental conditions and data comparisons performed in this study. (**A**) prior to and (**B)** during ATs infection. (**C**) Experimental strategy for RNA-seq and data analyses. Figure created using BioRender (https://biorender.com/).

Compared to unexposed *M.tb* (UE), A-ALF-exposed *M.tb* had 46 differentially expressed genes (DEGs) (29 upregulated and 17 downregulated) (A *vs*. UE, **Fig. 2A**), while E-ALF-exposed *vs.* unexposed *M.tb* had a total of 131 DEGs (116 upregulated and 15 downregulated) (E *vs*. UE, **Fig. 2A**). 23 DEGs were shared between the two (**Fig. 2B, Suppl. Fig. S2A**). DEGs were further sorted into functional categories (**Fig. 2C** and **D**, **Suppl. Table S1, Suppl. Table S2**). In A *vs*. UE, most DEGs associated with “Lipid metabolism” (6/7), “Conserved hypotheticals” (6/7), and “Information pathways” (1/1) were upregulated, whereas most DEGs linked to “Cell wall and cell processes” (5/7) and “Regulatory proteins” (2/3) were downregulated (**Fig. 2C**). Like A *vs*. UE, E *vs*. UE showed upregulation of most DEGs associated with “Conserved hypotheticals” (32/34), “Information pathways” (3/3), and “Lipid metabolism” (9/9), as well as downregulation of “Regulatory proteins” (3/4) (**Fig. 2D**). However, contrary to A *vs*. UE, in E *vs*. UE most “Cell wall and cell processes” (26/28) DEGs were upregulated, while “Insertion seqs and phages” (1/1) DEGs were downregulated (**Fig. 2D**).

**Figure 2.**
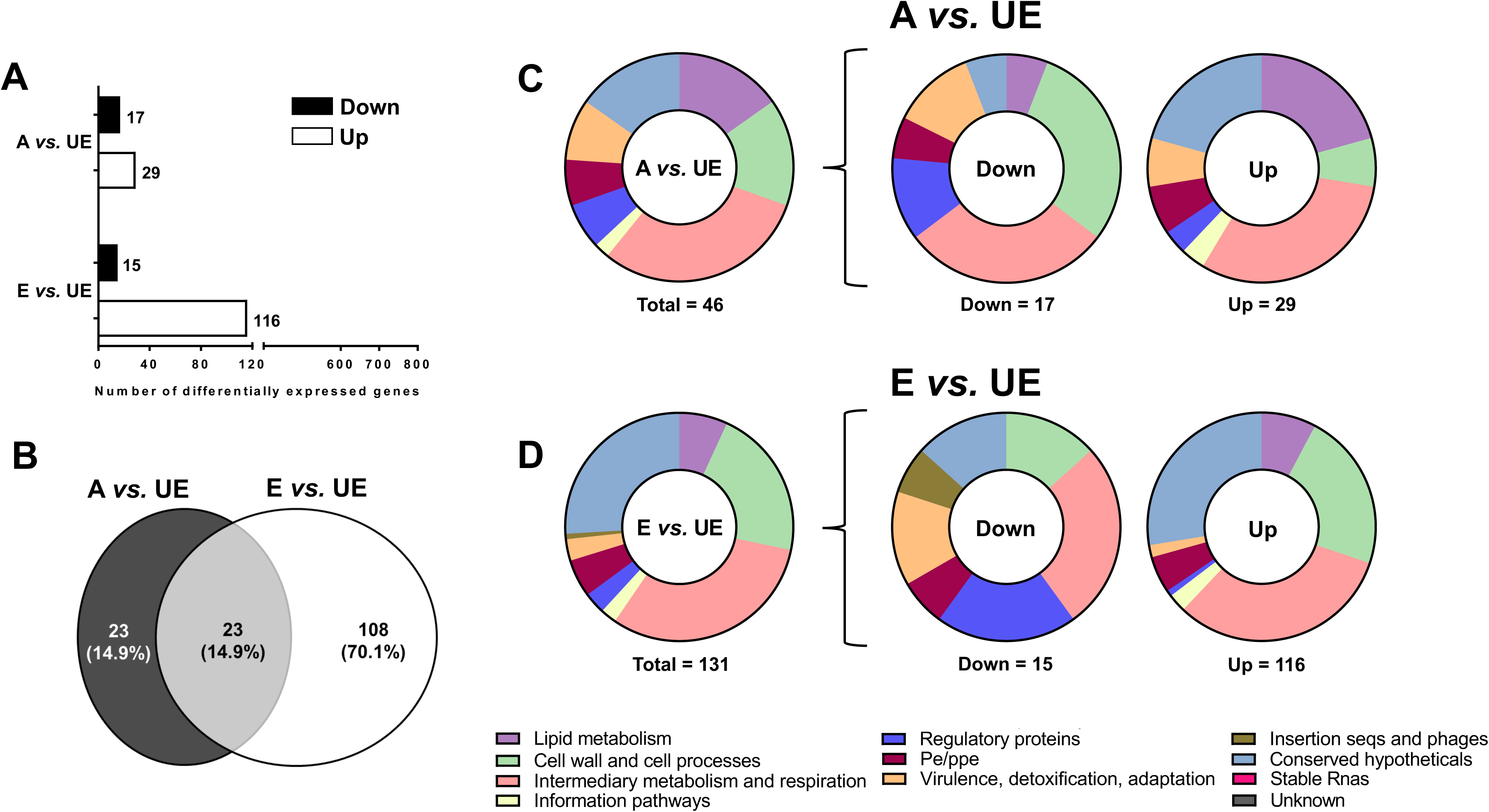
Differential expression analysis of ALF-exposed *M.tb* prior to AT infection. (**A**) Number of differentially expressed genes in ALF-exposed *M.tb vs.* unexposed *M.tb* prior to infection obtained with the DESeq2 method in the iDEP software, with the following settings: Log_2_ FC equal or greater than an absolute value of 1, FDR < 0.1. Upregulated genes are shown in white bars and downregulated genes in black bars. (**B**) Venn diagram of DEGs of ALF-exposed *M.tb vs*. unexposed *M.tb* prior to infection showing unique and shared DEGs in ALF-exposed *M.tb* prior to ATs infection. (**C**) Total number of DEGs distributed by Functional Categories for A-ALF-exposed *vs.* Unexposed *M.tb* (A *vs.* UE) and (**D**) for E-ALF exposed *vs.* Unexposed *M.tb* (E *vs.* UE) prior to ATs infection (represented as percentage). Graphs were generated in GraphPad Prism v9.1.1. A: A-ALF-exposed *M.tb*; E: E-ALF-exposed *M.tb*; UE: unexposed *M.tb*.

### E-ALF-exposed *M.tb* upregulates genes associated with ESX-secretion systems and phospholipases

*M.tb* has acquired specialized protein transport machinery including the type VII secretion systems (T7SSs) or ESX systems (25), composed of five distinct mycobacterial T7SSs (ESX-1 to ESX-5), with significant roles in virulence, such as nutrient uptake, immune evasion (*e.g*., phagosomal escape), immune modulation, horizontal gene transfer, and cell physiology (25, 26). Thus, we evaluated DEGs associated with all five ESX secretion systems (**Fig. 3**) as a potential mechanism associated with *M.tb* translocation into the cytosol (19, 27–29). Although expression of the majority of the genes related to the ESX-1, ESX-2, and ESX-3 secretion systems was not significantly altered in either A or E *vs*. UE, E *vs*. UE exhibited a higher (log_2_ FC >1) but not statistically significant (FDR >0.1) expression of ESX-1 genes *Rv3904C* (*esxE*) and *Rv3903C* (*cpnT*) (**Fig. 3A**), reported to mediate phagosomal disruption in *M.tb* (28, 30). In addition, phagosomal rupture and translocation of *Mycobacterium abscessus* into the host cytosol is facilitated by the ESX-4 secretion system (29), which substitutes for the ESX-1 secretion system in *M. abscessus* (29, 31). Interestingly, our results indicate that only the exposure to E-ALF resulted in a significant upregulation of DEGs (*eccC4*, *eccD4*, and *mycP4*) associated with the ESX-4 secretion system (**Fig. 3B**). While ESX-1, ESX-2, and ESX-4 secretion systems are essential for *M.tb* growth, the ESX-3 and ESX-5 have been related to immune modulation (26). In this regard, we found transcriptional changes in some PE/PPE genes associated with the ESX-5 system such as *PPE18* (significantly upregulated in A *vs*. UE) and *PPE65* (significantly upregulated in E *vs*. UE), and their functions are attributed to immunological regulation (32) (**Fig. 3C**).

**Figure 3.**
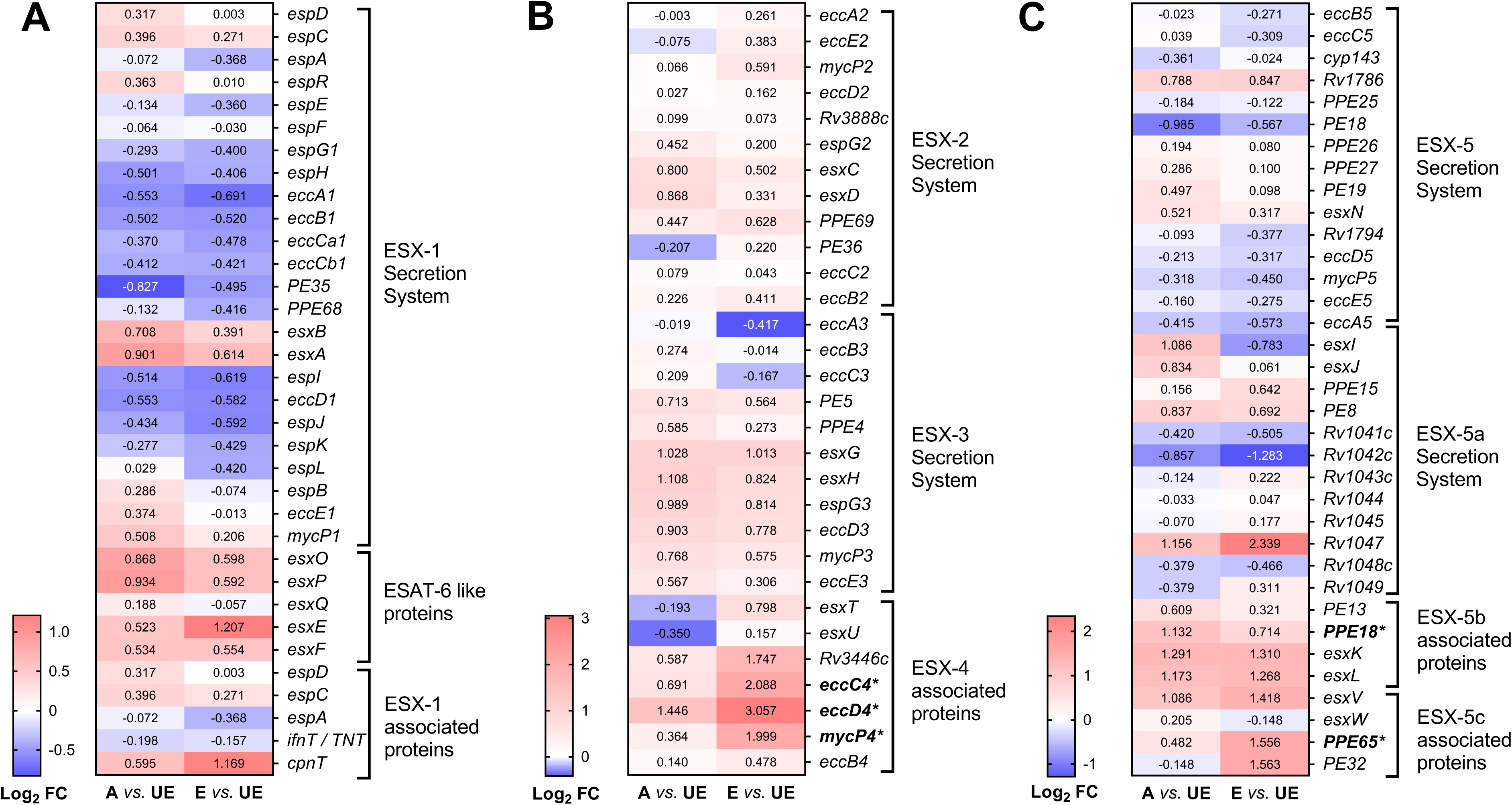
Heatmaps of *M.tb* genes associated with ESX-secretion systems prior to AT infection. (**A**) ESX-1 and ESAT-6 like proteins; (**B**) ESX-2, ESX-3, and ESX-4; and the (**C**) ESX-5 secretion system after A- or E-ALF exposure, prior to infection. Cells depict Log_2_ FC values in ALF-exposed *M.tb* (A: Adult ALF; E: Elderly ALF) *vs*. unexposed bacteria (UE), upregulated: red, downregulated: blue. Genes in bold indicate significant DEGs (Log_2_ FC equal or greater than an absolute value of 1, and FDR < 0.1) for both conditions. Genes with an asterisk indicate significance in only one of the comparisons, highlighting differences between conditions. Notice the different scales used in A, B, and C, for better visualization of the results. Heatmaps were generated in GraphPad Prism v9.1.1.

We also aimed to describe how exposure to human ALF modulates additional virulence factors involved in *M.tb* pathogenesis. Interestingly, like the production of phospholipases by *Listeria* to escape into the cytosol (33, 34), we found that phospholipase-related genes *Rv2349C* (*plcC*), *Rv2350C* (*plcB*), and *Rv2351C* (*plcA*) were uniquely upregulated in E *vs*. UE (**Table 1**). In addition, a few other virulence genes that encode phosphatases and proteases were only upregulated in E *vs*. UE, including *Rv2577*, *Rv1983* (*PE_PGRS35*), and *Rv3090* (**Table 1**). These are described to play a role in *M.tb* pathogenicity by promoting phagosomal arrest, increased replication, and dissemination (35, 36).

**Table 1.**
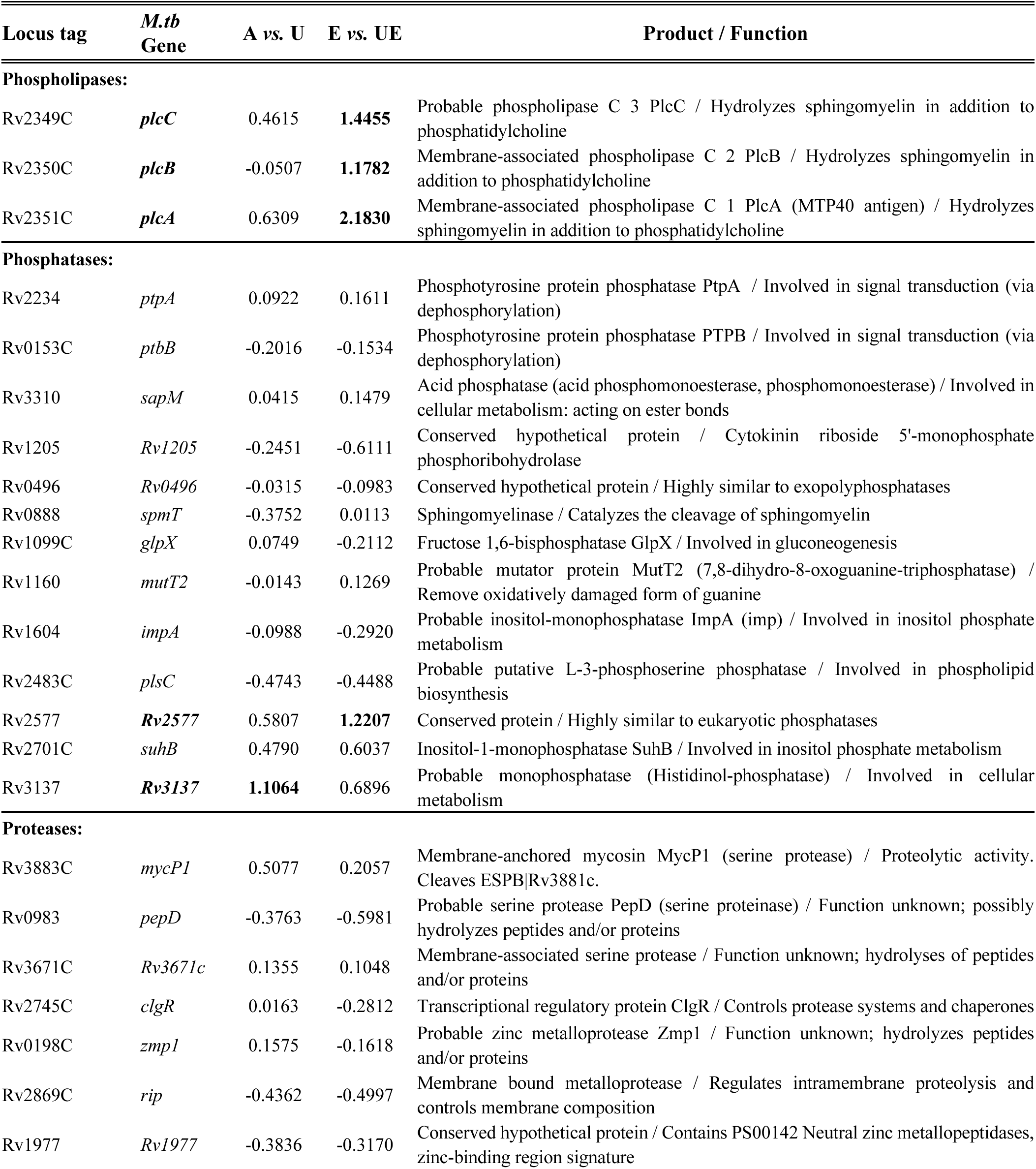
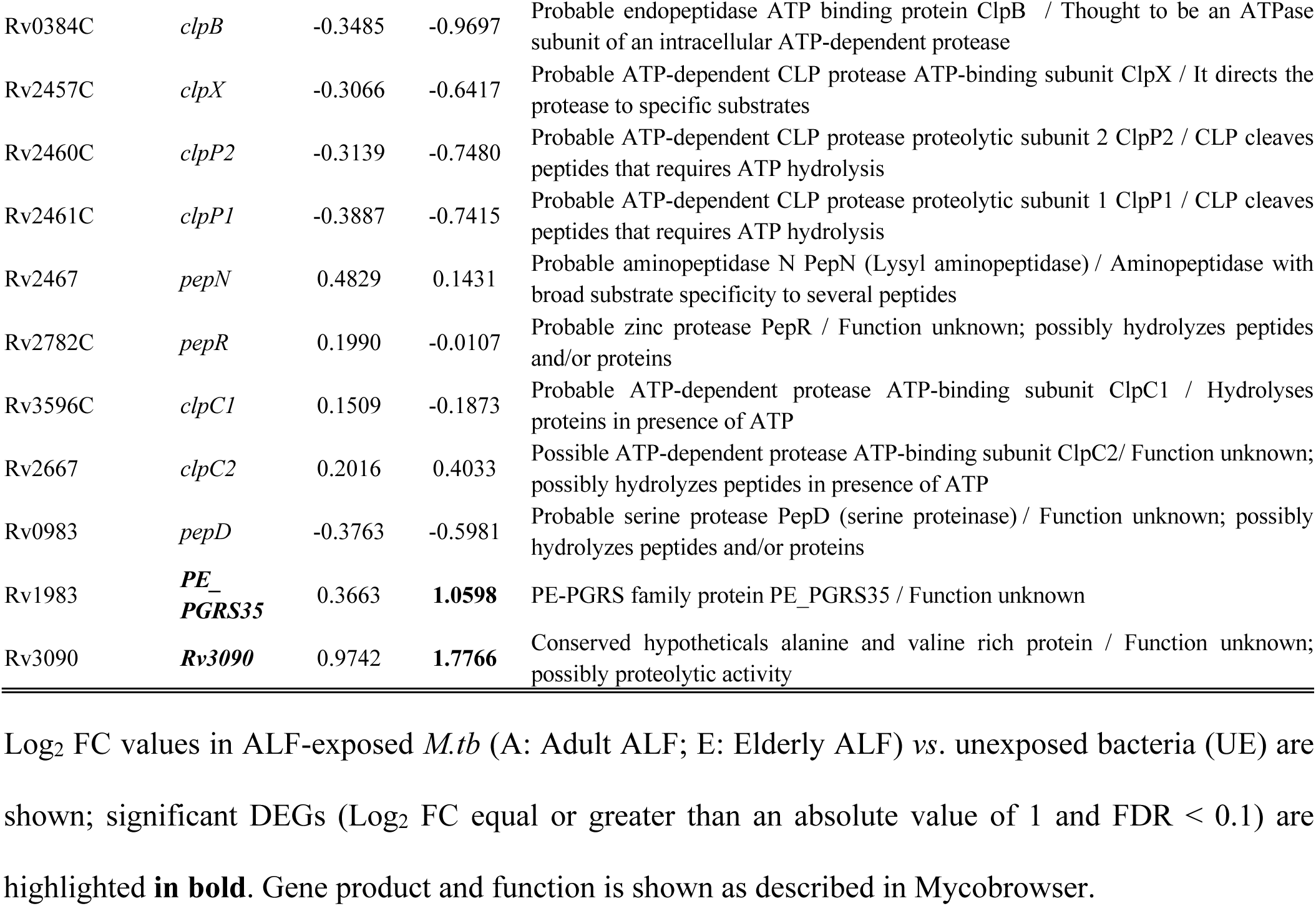
*M.tb* genes encoding phospholipases, phosphatases, and proteases after exposure to A-ALF or E-ALF before infection.

| Locus tag | *M.tb*<br>Gene | A vs. U | E vs. UE | Product / Function |
| --- | --- | --- | --- | --- |
| **Phospholipases:** |  |  |  |  |
| Rv2349C | *plcC* | 0.4615 | **1.4455** | Probable phospholipase C 3 PlcC / Hydrolyzes sphingomyelin in addition to phosphatidylcholine |
| Rv2350C | *plcB* | -0.0507 | **1.1782** | Membrane-associated phospholipase C 2 PlcB / Hydrolyzes sphingomyelin in addition to phosphatidylcholine |
| Rv2351C | *plcA* | 0.6309 | **2.1830** | Membrane-associated phospholipase C 1 PlcA (MTP40 antigen) / Hydrolyzes sphingomyelin in addition to phosphatidylcholine |
| **Phosphatases:** |  |  |  |  |
| Rv2234 | *ptpA* | 0.0922 | 0.1611 | Phosphotyrosine protein phosphatase PtpA / Involved in signal transduction (via dephosphorylation) |
| Rv0153C | *ptbB* | -0.2016 | -0.1534 | Phosphotyrosine protein phosphatase PTPB / Involved in signal transduction (via dephosphorylation) |
| Rv3310 | *sapM* | 0.0415 | 0.1479 | Acid phosphatase (acid phosphomonoesterase, phosphomonoesterase) / Involved in cellular metabolism: acting on ester bonds |
| Rv1205 | *Rv1205* | -0.2451 | -0.6111 | Conserved hypothetical protein / Cytokinin riboside 5'-monophosphate phosphoribohydrolase |
| Rv0496 | *Rv0496* | -0.0315 | -0.0983 | Conserved hypothetical protein / Highly similar to exopolysphosphatases |
| Rv0888 | *spmT* | -0.3752 | 0.0113 | Sphingomyelinase / Catalyzes the cleavage of sphingomyelin |
| Rv1099C | *glpX* | 0.0749 | -0.2112 | Fructose 1,6-bisphosphatase GlpX / Involved in gluconeogenesis |
| Rv1160 | *mutT2* | -0.0143 | 0.1269 | Probable mutator protein MutT2 (7,8-dihydro-8-oxoguanine-triphosphatase) / Remove oxidatively damaged form of guanine |
| Rv1604 | *impA* | -0.0988 | -0.2920 | Probable inositol-monophosphatase ImpA (imp) / Involved in inositol phosphate metabolism |
| Rv2483C | *plsC* | -0.4743 | -0.4488 | Probable putative L-3-phosphoserine phosphatase / Involved in phospholipid biosynthesis |
| Rv2577 | **Rv2577** | 0.5807 | **1.2207** | Conserved protein / Highly similar to eukaryotic phosphatases |
| Rv2701C | *suhB* | 0.4790 | 0.6037 | Inositol-1-monophosphatase SuhB / Involved in inositol phosphate metabolism |
| Rv3137 | **Rv3137** | **1.1064** | 0.6896 | Probable monophosphatase (Histidinol-phosphatase) / Involved in cellular metabolism |
| **Proteases:** |  |  |  |  |
| Rv3883C | *mycP1* | 0.5077 | 0.2057 | Membrane-anchored mycosin MycP1 (serine protease) / Proteolytic activity. Cleaves ESPB\|Rv3881c. |
| Rv0983 | *pepD* | -0.3763 | -0.5981 | Probable serine protease PepD (serine proteinase) / Function unknown; possibly hydrolyzes peptides and/or proteins |
| Rv3671C | *Rv3671c* | 0.1355 | 0.1048 | Membrane-associated serine protease / Function unknown; hydrolyses of peptides and/or proteins |
| Rv2745C | *clgR* | 0.0163 | -0.2812 | Transcriptional regulatory protein ClgR / Controls protease systems and chaperones |
| Rv0198C | *zmp1* | 0.1575 | -0.1618 | Probable zinc metalloprotease Zmp1 / Function unknown; hydrolyzes peptides and/or proteins |
| Rv2869C | *rip* | -0.4362 | -0.4997 | Membrane bound metalloprotease / Regulates intramembrane proteolysis and controls membrane composition |
| Rv1977 | *Rv1977* | -0.3836 | -0.3170 | Conserved hypothetical protein / Contains PS00142 Neutral zinc metalloproteases, zinc-binding region signature |
| Rv0384C | *clpB* | -0.3485 | -0.9697 | Probable endopeptidase ATP binding protein ClpB / Thought to be an ATPase subunit of an intracellular ATP-dependent protease |
| Rv2457C | *clpX* | -0.3066 | -0.6417 | Probable ATP-dependent CLP protease ATP-binding subunit ClpX / It directs the protease to specific substrates |
| Rv2460C | *clpP2* | -0.3139 | -0.7480 | Probable ATP-dependent CLP protease proteolytic subunit 2 ClpP2 / CLP cleaves peptides that requires ATP hydrolysis |
| Rv2461C | *clpP1* | -0.3887 | -0.7415 | Probable ATP-dependent CLP protease proteolytic subunit 1 ClpP1 / CLP cleaves peptides that requires ATP hydrolysis |
| Rv2467 | *pepN* | 0.4829 | 0.1431 | Probable aminopeptidase N PepN (Lysyl aminopeptidase) / Aminopeptidase with broad substrate specificity to several peptides |
| Rv2782C | *pepR* | 0.1990 | -0.0107 | Probable zinc protease PepR / Function unknown; possibly hydrolyzes peptides and/or proteins |
| Rv3596C | *clpC1* | 0.1509 | -0.1873 | Probable ATP-dependent protease ATP-binding subunit ClpC1 / Hydrolyses proteins in presence of ATP |
| Rv2667 | *clpC2* | 0.2016 | 0.4033 | Possible ATP-dependent protease ATP-binding subunit ClpC2/ Function unknown; possibly hydrolyzes peptides in presence of ATP |
| Rv0983 | *pepD* | -0.3763 | -0.5981 | Probable serine protease PepD (serine proteinase) / Function unknown; possibly hydrolyzes peptides and/or proteins |
| Rv1983 | **PE_**<br>**PGRS35** | 0.3663 | **1.0598** | PE-PGRS family protein PE_PGRS35 / Function unknown |
| Rv3090 | **Rv3090** | 0.9742 | **1.7766** | Conserved hypotheticals alanine and valine rich protein / Function unknown; possibly proteolytic activity |

Log<sub>2</sub> FC values in ALF-exposed *M.tb* (A: Adult ALF; E: Elderly ALF) vs. unexposed bacteria (UE) are shown; significant DEGs (Log<sub>2</sub> FC equal or greater than an absolute value of 1 and FDR < 0.1) are highlighted in **bold**. Gene product and function is shown as described in Mycobrowser.

### Differential expression analysis of ALF-exposed *M.tb* during AT infection

We reported that *M.tb* exposed to human E-ALF replicates faster in human macrophages and ATs *in vitro* and in the lungs of *mice in vivo* (13, 15). Further, in ATs, we found that at 72 h, *M.tb* translocate from endosomal compartments into the cytosol, conferring an advantageous niche for faster *M.tb* replication and survival within ATs (15). Thus, herein we determined the transcriptional changes of *M.tb* exposed to A-ALF (A_i_) or E-ALF (E_i_) during AT infection compared to ALF-exposed *M.tb* prior to infection (A or E) (A_i_ *vs*. A, and E_i_ *vs*. E comparisons, respectively, **Fig. 4A**). According to the DE analysis, both comparisons had similar number of upregulated and downregulated genes (**Fig 4A**) and shared 1,059 DEGs (**Fig. 4B, Suppl. Fig. S2B**). Comparable numbers of DEGs in each functional category were found in both (**Fig. 4C** and **D, Suppl. Table S1, Suppl. Table S2**). Therefore, the functional category distribution did not differ remarkably between A_i_ *vs*. A, and E_i_ *vs*. E (**Fig. 4C** and **D**).

**Figure 4.**
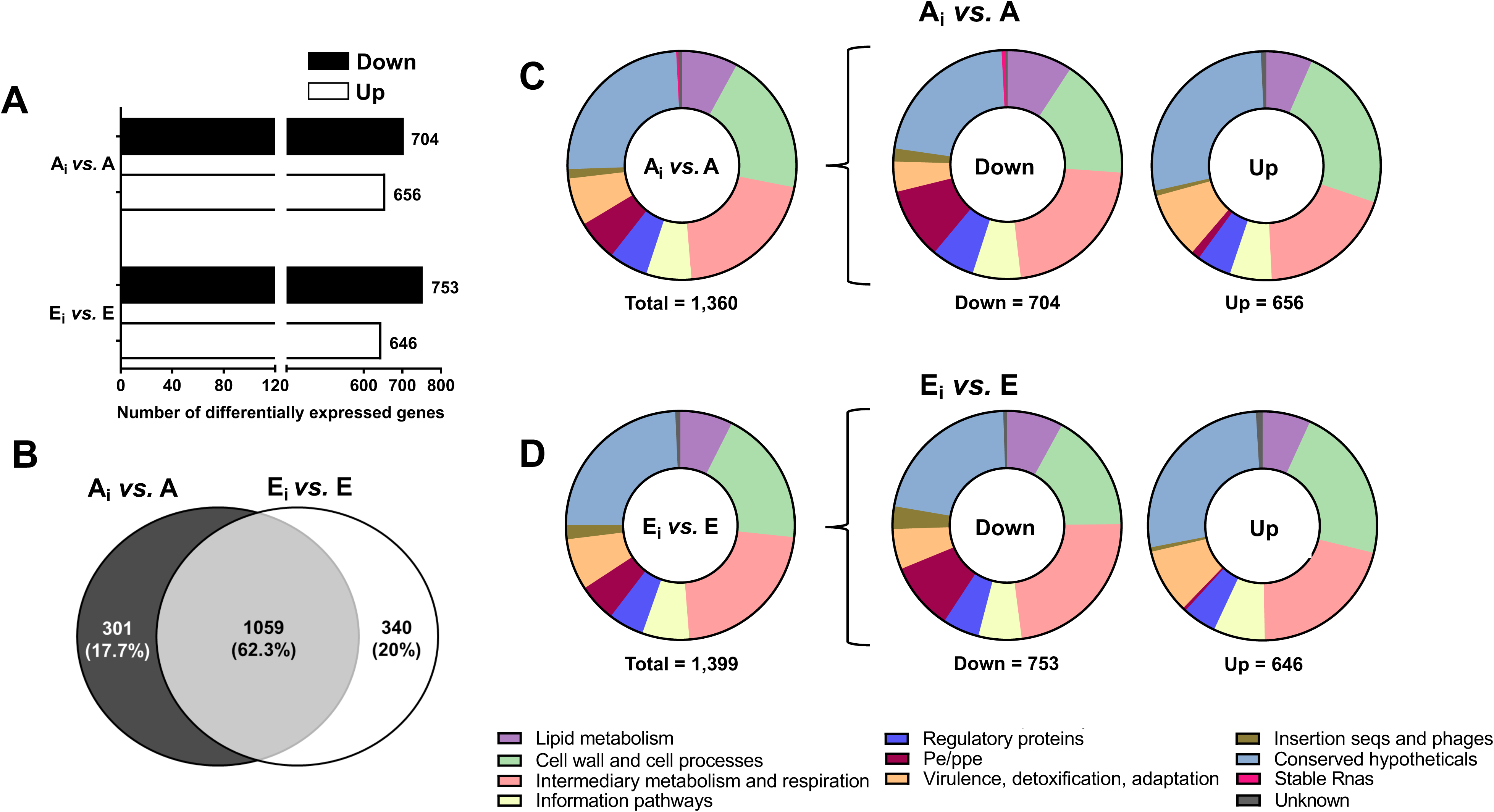
Differential expression analysis of ALF-exposed *M.tb* during AT infection. (**A**) Number of differentially expressed genes during AT infection (ALF-exposed *M.tb* in ATs *vs*. ALF-exposed *M.tb* before infection) obtained with the DESeq2 method in the iDEP software, with the following settings: Log_2_ FC equal or greater than an absolute value of 1, and FDR < 0.1. Upregulated genes are shown in white bars and downregulated genes in black bars. (**B**) Venn diagram of DEGs during AT infection (ALF-exposed *M.tb* in ATs *vs*. ALF-exposed *M.tb* before infection) showing unique and shared DEGs in ALF-exposed *M.tb* during AT infection. (**C**) Total number of DEGs distributed by Functional Categories during AT infection for A-ALF-exposed *M.tb* in ATs *vs*. A-ALF-exposed *M.tb* prior infection (A_i_ *vs.* A) and (**D**) for E-ALF-exposed *M.tb* in ATs *vs*. E-ALF-exposed *M.tb* prior infection (E_i_ *vs.* E) during ATs infection (represented as percentage). Graphs were generated in GraphPad Prism v9.1.1. A: A-ALF-exposed *M.tb*; E: E-ALF-exposed *M.tb*; A_i_: A-ALF-exposed *M.tb* during AT infection; E_i_: E-ALF-exposed *M.tb* during AT infection.

### Effects of A- *vs.* E-ALF exposure on the expression of *M.tb* ESX-associated genes during AT infection

Our results indicate that when *M.tb* is exposed to either A-ALF or E-ALF, most of the ESX-1-related genes, including those encoding for ESAT-6 (*esxA*) and CFP-10 (*esxB*), become downregulated during infection in ATs (A_i_ *vs*. A, and E_i_ *vs*. E, **Fig. 5A**). This includes the ESX-1-associated gene *espI* (*Rv3876*) that mediates ESAT-6 secretion (27), which was downregulated in both comparisons but was only significant in A _i_ *vs*. A (**Fig. 5A**). Conversely, *ifnT/TNT* was the only ESX-1-associated gene that was upregulated in both comparisons. The only ESX-2-related gene that was differentially expressed and downregulated in A_i_ *vs*. A was *espG2*, with unknown function (**Fig. 5B**). Further, most of the ESX-3 genes (*e.g.*, *PE5*, *esxH*, *espG3*, *mycP3*, and *eccE3*) were downregulated in both comparisons, except for *esxG* (*Rv0287*), involved in iron regulation, which was significantly downregulated only in E_i_ *vs*. E, as well as ESX-4-associated genes *Rv3446c* and *eccC4* (**Fig. 5B**). Interestingly, related to the immunomodulatory ESX-5 secretion system, two CFP-10 homologs playing crucial roles in regulating cytokine/chemokine signaling (31, 32), *esxJ* (*Rv1038C*) and *esxW* (*Rv3620c*), were found significantly upregulated in E_i_ *vs*. E (*esxJ*) and downregulated in A_i_ *vs*. A (*esxW*) (**Fig. 5C**). In addition, *PPE26* and *PPE65,* involved in pro-inflammatory responses (32, 37), were significantly downregulated in E_i_ *vs*. E, but upregulated in A_i_ *vs*. A (**Fig. 5C**). Other ESX-5-associated genes (*PPE27*, *esxI*, *Rv1042c*, *Rv1048c*, and *esxV*) were upregulated in both A_i_ *vs*. A, and E_i_ *vs*. E (**Fig. 5C**).

**Figure 5.**
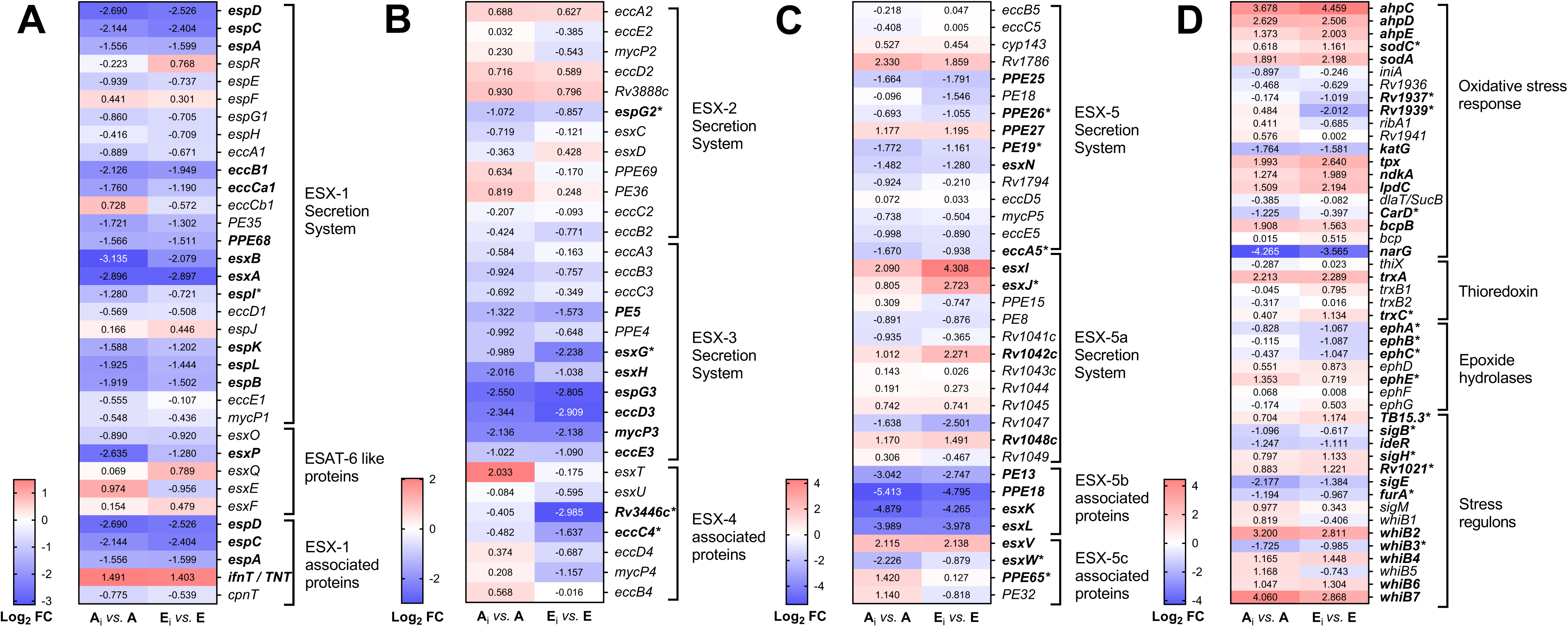
*M.tb* genes associated with ESX-secretion systems and ROS defense mechanisms at 72 hpi in ATs. (**A**) ESX-1 and ESAT-6 like proteins; (**B**) ESX-2, ESX-3, and ESX-4 secretion systems; (**C**) ESX-5 secretion system, and (**D**) ROS defense mechanisms, after infection of ATs with A-ALF or E-ALF-exposed *M.tb* for 72 hours. Heatmap cells depict Log_2_ FC values in ALF-exposed *M.tb* in ATs (A_i_: Adult ALF-exposed *M.tb* during infection; E_i_: Elderly ALF-exposed during infection) *vs*. ALF-exposed *M.tb* before infection (A: Adult ALF; E: Elderly ALF), upregulated: red, downregulated: blue. Genes in bold indicate significant DEGs (Log_2_ FC equal greater than an absolute value of 1, and FDR < 0.1) for both conditions. Genes with an asterisk indicate significance in only one of the comparisons, highlighting differences between conditions. Notice the different scales used in A, B, C, and D for better visualization of the results. Heatmaps were generated in GraphPad Prism v9.1.1.

### Effects of A- *vs.* E-ALF exposure on the expression of PDIM-related genes during *M.tb* replication in ATs

Most *ppsA-E* genes involved in producing phthiocerol and phenolphthiocerol (38) were significantly downregulated in *M.tb* exposed to A-ALF during ATs infection (**Table 2**), except for *ppsC* that was also significantly downregulated in E-ALF-exposed bacteria. Although not significant, the transcriptional repressor, *Rv3167c*, which controls the lipid PDIM operon (23), was downregulated in E-ALF-exposed bacteria during AT infection (E_i_ *vs*. E, **Table 2**). Further, in both comparisons (E_i_ *vs*. E and A_i_ *vs*. A) most of the polyketide synthases-related genes (*pks1*, *pks10*, *pks12*, *pks5,* and *pks7*), involved in PDIM biosynthesis (39), were downregulated (**Table 2**), as well as MmpL7 (40), involved in PDIMs and phenolic glycolipid (PGL) transport across the membrane. In contrast, other PDIMs and antibiotic-related transporters were found upregulated [*e.g.*, *drrB* and *drrC* (41, 42)] (**Table 2**).

**Table 2.**
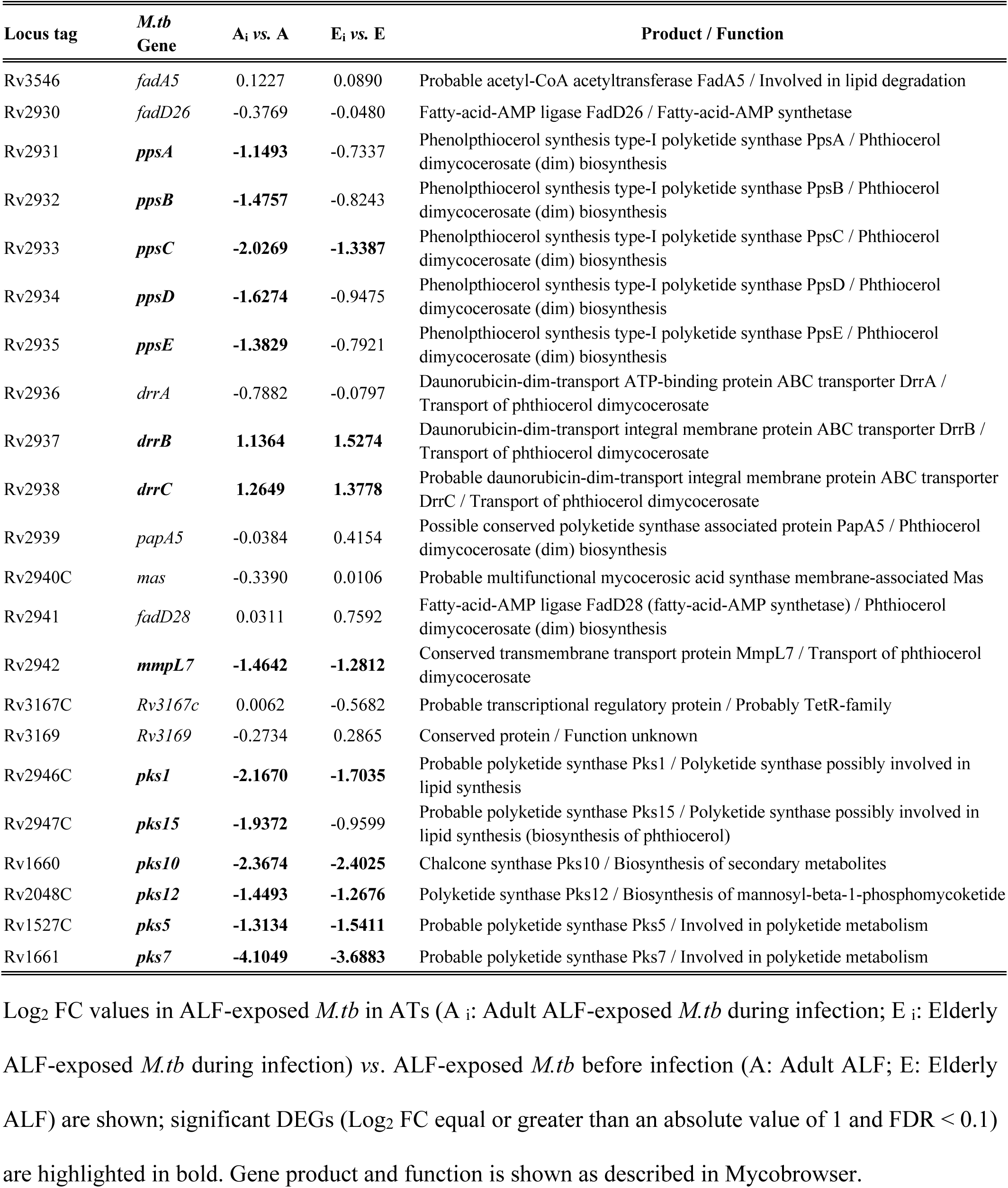
*M.tb* genes related to lipid phthiocerol dimycoserosate (PDIMs) biosynthesis and translocation during infection in ATs.

### E-ALF-exposed *M.tb* has upregulation of genes associated with the Sec (secretion) system during AT infection

Protein transport through the *M.tb* cell membrane is mainly mediated by the general secretion (Sec) pathway and the twin-arginine translocation (Tat) system, in addition to the ESX-secretion system (43–45). We assessed DEGs linked to these two *M.tb* export systems in both comparisons (E_i_ *vs*. E and A_i_ *vs*. A) (**Table 3**). Interestingly, E_i_ *vs*. E showed upregulation of *secA1*, *secE1*, and *yajC* genes during infection, the latter encoding an accessory Sec protein that aids in protein export (43) (**Table 3**). Further, *M.tb* uses the signal recognition particle system (SRP pathway) to secrete co-translational proteins from ribosomes to the bacterial extracellular surface (46). The SRP pathway comprises three major components, FtsY, FfH, and 4.5S rRNA (46, 47). In this regard, E_i_ *vs*. E had upregulation of the *ffh* gene in ATs (**Table 3**). Lastly, we found upregulation of the *tatA* gene only in the A_i_ *vs*. A comparison (**Table 3**), which encodes for one of the three integral membrane proteins (TatA-C) of the twin-arginine translocation system, important to export of folded proteins across the bacterial cell envelope (45).

**Table 3.** *M.tb* genes associated with the Sec secretion and Tat translocation systems during replication in Ats.

| Locus tag | *M.tb* Gene | A<sub>i</sub> vs. A | E<sub>i</sub> vs. E | Product / Function |
| --- | --- | --- | --- | --- |
| **Sec Secretion System:** |  |  |  |  |
| Rv3240C | *secA1* | 0.5768 | **1.2521** | Probable preprotein translocase SecA1 1 subunit / Involved in protein export |
| Rv1821 | *secA2* | -0.4800 | 0.1331 | Possible preprotein translocase ATPase SecA2 /Involved in protein export |
| Rv2587C | *secD* | -0.1348 | 0.0324 | Probable protein-export membrane protein SecD |
| Rv0638 | *secE1* | 0.6131 | **1.1954** | Probable preprotein translocase SecE1 / Essential for protein export |
| Rv2586C | *secF* | -0.5924 | -0.3857 | Probable protein-export membrane protein SecF / Involved in protein export |
| Rv1440 | *secG* | **1.4243** | **1.9535** | Probable protein-export membrane protein (translocase subunit) SecG/ Participates in an early event of protein translocation |
| Rv0732 | *secY* | **2.0271** | **2.6184** | Probable preprotein translocase SecY / Essential for protein export |
| Rv2916C | *ffh* | 0.5952 | **1.0993** | Probable signal recognition particle protein Ffh / Necessary for efficient export of extra-cytoplasmic proteins |
| Rv2921C | *ftsY* | -0.1439 | 0.3242 | Probable cell division protein FtsY (SRP receptor) / Reception and insertion of a subset of proteins at the membrane |
| Rv2588C | *yajC* | 0.7343 | **1.3427** | Probable conserved membrane protein secretion factor YajC / Involved in secretion apparatus |
| Rv3921C | *yidC* | 0.6739 | 0.7670 | Probable conserved transmembrane protein / Cell wall and cell processes |
| **Tat Secretion System:** |  |  |  |  |
| Rv2094C | *tatA* | **1.4052** | 0.9527 | Sec-independent protein translocase membrane-bound protein TatA / Involved in protein export |
| Rv1224 | *tatB* | 0.6343 | 0.6229 | Probable protein TatB / Twin-arginine translocation (Tat) system |
| Rv2093C | *tatC* | 0.5748 | 0.0482 | Sec-independent protein translocase transmembrane protein TatC / Involved in protein export |

Log<sub>2</sub> FC values in ALF-exposed *M.tb* in ATs (A<sub>i</sub>: Adult ALF-exposed *M.tb* during infection; E<sub>i</sub>: Elderly ALF-exposed *M.tb* during infection) vs. ALF-exposed *M.tb* before infection (A: Adult ALF; E: Elderly ALF) are shown; significant DEGs (Log<sub>2</sub> FC equal or greater than an absolute value of 1 and FDR < 0.1) are highlighted in bold. Gene product and function is shown as described in Mycobrowser.

### Upregulation of E-ALF-exposed *M.tb* genes encoding oxidative stress defense mechanisms during AT infection

*M.tb* has developed strategies that mitigate oxidative stress and overcome the microbicidal capacity within the host (48, 49). We next explored DEGs linked to reactive oxygen species (ROS) defense mechanisms (**Fig. 5D; Suppl. Fig. S3)**. While *ahpC*, *ahpD*, *ahpE*, *lpdC*, *trxA*, and *sodA* genes were upregulated in both comparisons (E_i_ *vs*. E and A_i_ *vs*. A), *sodC* and *trxC* were significantly upregulated only in the E_i_ *vs*. E (**Fig. 5D**), all associated with response to oxidative stress and defense against the microbicidal capacity of the host (50).

In addition, epoxide hydrolases, sigma factors, and WhiB proteins are all characterized as additional *M.tb* detoxifying strategies to withstand adverse conditions (51–53). Our findings indicate that during infection E-ALF-exposed *M.tb* downregulates some epoxide hydrolases genes (*ephA*, *ephB,* and *ephC*) (**Fig. 5D**). Conversely, sigma factor *sigH*, which is considered a crucial *M.tb* transcriptional factor responding to oxidative, nitrosative, and heat stressors (52, 54), was found upregulated in E_i_ *vs*. E (**Fig. 5D**), while sigma factor B (*sigB*), which plays comparable roles (55), was downregulated in A_i_ *vs*. A (**Fig. 5D**). Finally, the WhiB family are redox-sensing transcription factors in which the three members (WhiB3, WhiB4, and WhiB7) are responsible in maintaining redox homeostasis (53). Our results indicate that *whiB2, whiB4, whiB6* and *whiB7* were upregulated in both comparisons, whereas *whiB3* was downregulated only in A_i_ *vs*. A (**Fig. 5D**). Other stress response-related genes were uniquely upregulated (E_i_ *vs*. E: *TB15*.3 and *Rv1021*; A_i_ *vs*. A: *ephE*) or downregulated (E_i_ *vs*. E: *Rv1937* and *Rv1939*; A_i_ *vs*. A: *furA* and *CarD*) (**Fig. 5D**), while others were shared between both comparisons (upregulated: *tpx* and *bcpB*; downregulated: *katG*, *narG*, *ideR* and *sigE*) (**Fig. 5D**).

### Upregulation of other virulent factors in E-ALF-exposed *M.tb* during AT infection

Additional virulence determinants that may contribute to *M.tb* pathogenesis including proteases (*zmp1*, *clpP1* and *pepN*) (56, 57), lipases (*lipE*) (58), iron efflux pumps (*MmpL4*) (59) and adhesins (*apa*, *fbpB*, *glnA1*, *gap* and *pstS1*) (60) were found upregulated in E_i_ *vs*. E, as well as other genes associated with mycobacterial survival (*glpX* and *Rv1096*) (61, 62) (**Suppl. Fig. S3**). Other upregulated or downregulated DEGs in E_i_ *vs.* E or A_i_ *vs.* A, or in both comparisons are depicted in **Suppl. Fig. S3**.

## DISCUSSION

The elderly population is considered a large reservoir for *M.tb* infection due to their increased vulnerability to active TB disease or reactivation of latent TB (7, 8). We have published that the lung mucosa or ALF from elderly individuals (65+ years old) is highly oxidized and constitutes a pro-inflammatory environment with dysfunctional innate proteins (12–14), and that *M.tb* replicates faster upon contacting the elderly lung mucosa (E-ALF) in human macrophages and can replicate to a higher bacterial load in mice, leading to more extensive lung tissue damage (13). More recently, we reported that *M.tb* exposure to E-ALF enhances intracellular growth in ATs and alters endosomal trafficking with increased bacterial translocation into the cytosol (15). Our findings herein suggest that age-associated ALF changes lead to an increased predisposition of the elderly population to *M.tb* infection. Thus, it is critical to understand how exposure to human ALF from adult or elderly individuals changes the *M.tb* metabolic status. In this study we investigated the transcriptional profile of *M.tb* when exposed to human ALF (A-ALF or E-ALF) before and during intracellular replication in ATs. Our DEG analysis revealed upregulation of genes associated with the ESX-secretion systems and phospholipases in E-ALF-exposed *M.tb* compared to unexposed *M.tb* before infection, but not in A-ALF-exposed *M.tb* (**Fig. 6**). During AT infection, E-ALF-exposed *M.tb* also showed altered expression of virulence-associated genes related to the ESX-secretion systems, as well as several genes associated with PDIMs metabolism and ROS defense strategies, among others (**Fig. 6**). Our findings suggest that *M.tb* takes advantage of E-ALF exposure by driving metabolic changes to avoid the intrinsic antimicrobial mechanisms of the host with potential implications in infection outcome.

**Figure 6.**
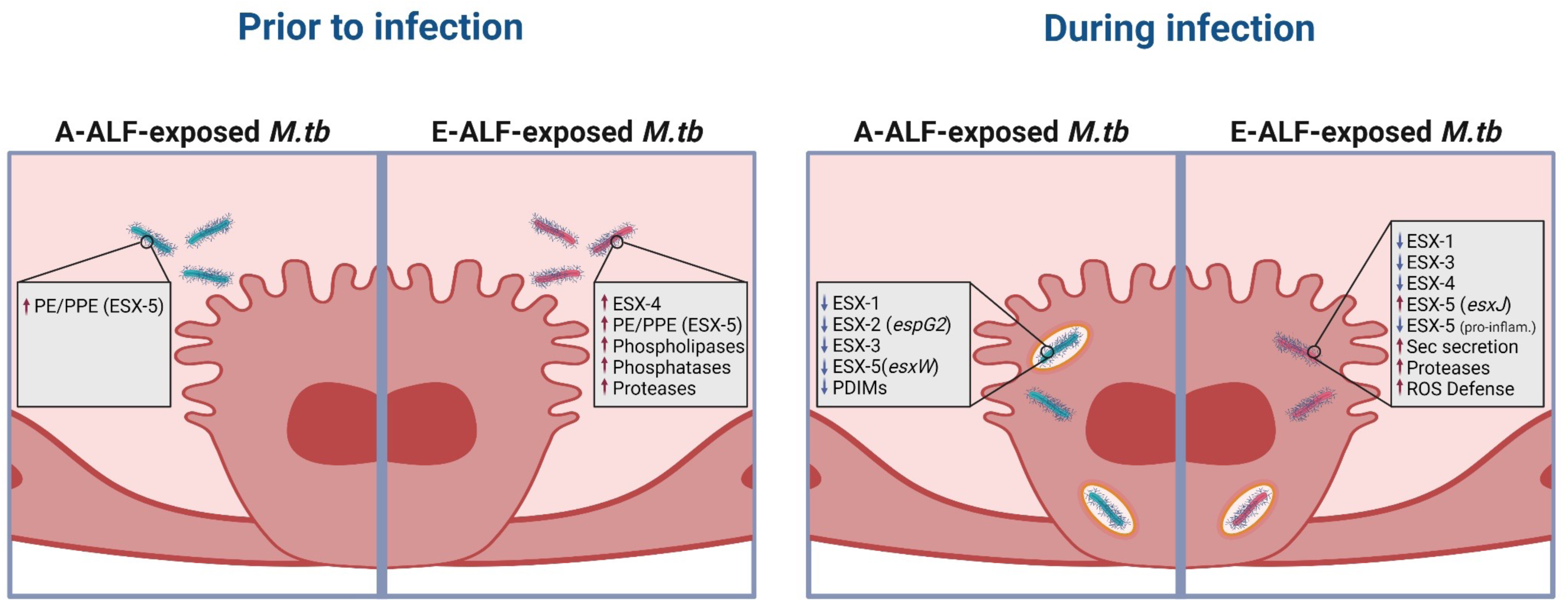
Schematic overview of the main findings in this study. Transcriptional profile of ALF-exposed-*M.tb* prior to and during intracellular replication in ATs. Figure created using BioRender (https://biorender.com/).

Indeed, *M.tb* has evolved a vast range of virulence factors implicated in lipid metabolism and protein transport, signal transduction, nutrient uptake, immune evasion, and resistance to microbicidal activity, among others (48, 63). For instance, *M.tb* employs five ESX protein secretion systems (ESX-1 to ESX-5). In general, full *M.tb* virulence requires the presence of ESX-1, ESX-3, and ESX-5 (25). However, the ESX-1 secretion system is considered one of the key virulence factors due to the secretion of the 6-kDa Early Secreted Antigenic Target (ESAT-6/EsxA) and its partner protein, the 10-kDa Culture Filtrate Protein (CFP-10/EsxB), associated with phagosomal rupture and bacterial translocation into the host cytosol (20, 25, 64).

In this regard, for *M.tb* exposure to ALF prior to infection, our results indicate that although the genes encoding ESAT-6 (*esxA*) and CFP-10 (*esxB*) are not significantly upregulated after exposure to either A- or E-ALF, *esxE* and *cpnT* are slightly more upregulated in the E-ALF group. Both genes are encoded by the *cpnT* operon, which may allow TB necrotizing toxin (TNT) secretion (CpnT/TNT) to contribute to the permeabilization of the phagosomal membrane (28, 30). Particularly, the EsxF/EsxE heterodimer generates persistent membrane-spanning channels that exhibit pore-forming activity (65). This highlights how the ESX family in *M.tb* has paralogous gene pairs, in which proteins encoded frequently form a 1:1 heterodimer (*e.g.*, EsxA-EsxB) (66) and cause lysis of the phagosome and bacterial translocation into the cytosol (19, 20). However, *M. abscessus* lacks the ESX-1 secretion system, but instead, has the ESX-4 secretion system with a similar role (29, 31). Interestingly, E-ALF-exposed *M.tb* (prior to infection) shows upregulation of genes associated with the ESX-4 secretion system (*Rv3446c*, *eccC4*, *eccD4*, and *mycP4*). Other ESX-5-related genes are differentially expressed, such as *PPE18* (upregulated in A-ALF-exposed *M.tb*) and *PPE65* (upregulated in E-ALF-exposed *M.tb*) involved in anti-inflammatory and pro-inflammatory signaling, respectively (32).

In addition, all phospholipase-related genes (*plcC*, *plcB,* and *plcA*), which mediate hydrolytic reactions with cytotoxic effects on macrophages (67), were upregulated when *M.tb* was exposed to the E-ALF but not A-ALF (prior to infection). Interestingly, the phosphatidylinositol-specific phospholipase C (PI-PLC/*plcA*) and the broad-range phospholipase C (PC-PLC/ *plcB*) enzymes, together with the pore-forming toxin listeriolysin O (LLO), are used by the intracellular pathogen *Listeria* to break the phagosomal membrane (33, 34). In *M.tb*, these virulence phospholipase homologs confer higher resistance to alveolar macrophage microbicidal activity (67) and are suggested to participate in the breach of vacuolar membranes (68). Additionally, some virulent-related phosphatases (*e.g. Rv2577*) and protease genes (*e.g. PE_PGRS35* and *Rv3090*) were also upregulated in the E *vs.* UE comparison. *Rv2577* is characterized as an alkaline phosphatase/phosphodiesterase enzyme, which plays a role in *M.tb* pathogenicity by promoting increased replication and phagosomal arrest in early endosomes (35), and *Rv3090* induces late cell apoptosis, resulting in bacterial dissemination and organ damage in mice (36). Overall, these results highlight how exposure to the human elderly lung mucosa prior to infection drives the upregulation of virulence-related genes in *M.tb*, potentially priming the bacterium to upregulate specific functions involved in mitigating the host immune response and ready its machinery to permeabilize phagosomes before contacting host cells.

We next investigated the effects of E-ALF exposure on *M.tb* metabolism during infection of ATs. In contrast with previous studies that reported upregulation of ESX-1-associated genes (69), our results indicate that most ESX-1 genes are downregulated after exposure to ALF (either A- or E-ALF) during AT infection. Even though the expression of *esxA* is not upregulated, its levels could be influenced by additional ESX proteins that mediate its secretion. For instance, A-ALF-exposed *M.tb* had downregulation of *EspI,* which regulates ESAT-6 secretion, impacting its availability and potentially levels of phagosomal disruption.

However, our findings revealed upregulation of some DEGs associated with ESX-5 during AT infection. Interestingly, *esxJ,* homolog of CFP-10/*esxB*, was significantly upregulated in E-ALF-exposed *M.tb* during infection, while the other homolog *esxW* was significantly downregulated in A-ALF-but not in E-ALF-exposed *M.tb*. Further studies are needed to verify whether these homologs, mainly *esxJ*, may play a similar role to *esxB* from the ESX-1 system, leading to bacterial escape into the cytosol. ESX-5 immunomodulatory proteins, PPE26 and PPE65, responsible for pro-inflammatory responses in macrophages (32, 37), were both downregulated in E-ALF-exposed *M.tb* during replication in ATs, suggesting potential mechanisms of *M.tb* to avoid pro-inflammatory responses within the host leading to increased intracellular survival, consistent with our prior findings (15).

One of the main *M.tb* virulence factors are PDIMs, which play a broad range of roles during infection (70–72). Recent evidence suggests that PDIMs cooperate with *esxA* creating pores in the phagosomal membrane to allow access of *M.tb* and secreted proteins into the host cell cytosol, ultimately leading to cell apoptosis (23, 63, 72). In our dataset, A-ALF-exposed *M.tb* during infection in ATs had significant downregulation of genes encoding the phenolpthiocerol synthesis type-I polyketide synthase (*ppsA*, *ppsB*, *ppsC*, *ppsD,* and *ppsE*) that mediate PDIMs biosynthesis, while E-ALF-exposed *M.tb* only showed significant downregulation of *ppsC*. Polyketide synthases *pks1*, *pks10*, *pks12*, *pks5*, and *pks7*, involved in the production of PDIMs, were downregulated in both A- and E-ALF-exposed *M.tb* during infection. Conversely, PDIM transport genes *drrB* and *drrC* were the only genes significantly upregulated in both groups. Finally, the transcriptional repressor *Rv3167c*, which regulates the PDIM operon, was marginally downregulated in E-ALF-exposed bacteria. Interestingly, the loss-of-function of *Rv3167c* correlates with the capacity of *M.tb* to escape the phagosome (23).

In addition, mycobacteria use a wide range of secretion systems including the general secretion (Sec) pathway (SecA1 secretory pathway and the alternative SecA2-operated pathway) and a twin-arginine translocation system (Tat system) (43–45, 73). Our results indicate that *secA1*, essential for *M.tb* growth and responsible for exporting most proteins through the SecYEG complex (43, 44), and *secE1* genes are significantly upregulated during infection of ATs with E-ALF-exposed *M.tb* but not in A-ALF-exposed *M.tb*, whereas *secG* and *secY* genes are upregulated regardless of the ALF exposure condition. The *yajC* gene, which encodes an auxiliary Sec protein (43), was also upregulated only in E-ALF-exposed *M.tb*. Regarding the Tat system, we observed upregulation of the *tatA* gene during infection of ATs by A-ALF-exposed bacteria, which encodes for one of its integral membrane proteins (TatA-C) (45). Overall, these results suggest an increase in protein transport in intracellular *M.tb* that has been previously exposed to E-ALF when compared to A-ALF-exposed *M.tb*.

Further, mycobacteria have natural mechanisms to combat the microbicidal capacity of host cells, such as the production of virulence factors against oxidative and nitrosative stresses (48, 49). Indeed, *M.tb* genes encoding catalase-peroxidase (*katG*), superoxide dismutase (*sodA*), alkyl peroxide reductase (*ahpCDE*), thioredoxin (*thiX* and *trxB1C*), or thioredoxin reductase (*trxB2*) are upregulated in response to nitrosative and oxidative stressors (50). Several genes that encode ROS defense mechanisms were upregulated in both A-ALF and E-ALF-exposed *M.tb* during AT infection (e.g. *ahpC*, *ahpD*, *ahpE*, *trxA*, *lpdC*, and *sodA*). However, *sodC* and *trxC* were upregulated only in E-ALF-exposed *M.tb*. Superoxide dismutase (SOD) is a radical scavenger that reduces NO and oxygen radicals’ production, allowing *M.tb* to tolerate oxidative stress and survival within host cells (74), whereas thioredoxin (Trx) is involved in the maintenance of cellular redox equilibrium, contributing to *M.tb* resistance to phagocytosis (75). Additionally, sigma factors *sigH* and *sigB*, essential for bacterial adaptation to oxidative, nitric oxide, and heat stress (76, 77), were significantly upregulated in E-ALF-exposed *M.tb* but downregulated in A-ALF-exposed *M.tb*. Further, WhiB proteins perform a variety of functions including regulating redox homeostasis. Although *whiB2*, *whiB4*, *whiB6* and *whiB7* were upregulated in both A-ALF and E-ALF-exposed *M.tb* during infection of ATs, *whiB3*, necessary to mitigate redox stress caused by ROS and RNS and to survive within macrophages (53), was downregulated solely in the A-ALF condition.

Other virulence factors are required for *M.tb* growth and pathogenesis, such as proteases, lipases, and adhesins. The proteases *zmp1* and *clpP1,* and the lipase *lipE,* were upregulated in E-ALF-exposed *M.tb* during AT infection. Metalloprotease Zmp-1 is involved in inflammasome inactivation and phagosome maturation arrest and is essential for intracellular survival (78, 79), and clpP1 is critical for *M.tb* growth (80). Similarly, LipE with lipase/esterase activity is induced under stress conditions and is essential for *M.tb* intracellular growth *in vivo* (58). Lastly, several adhesins such as *apa*, *fbpB*, *glnA1*, *gap,* and *pstS1* were upregulated in E-ALF-exposed *M.tb* during infection. These adhesins are associated with adherence-mediating elements and strategies that permit *M.tb* attachment to the host (60).

Finally, a potential limitation of the study is the use of A- and E-ALF pools instead of individual ALFs. We chose this strategy to reduce human-to-human variability within each ALF group studied and focus on the overall effects of A- and E-ALF exposure rather than donor-specific changes. However, we understand that human variability strongly influences the outcomes of *M.tb* infection and thus, the impact of individual A- and E-ALFs on *M.tb* adaptation to the lung environment will be investigated in further studies. In addition, we note here that for one of the experimental conditions (E-ALF-exposed *M.tb* before infection or E-ALF) we only have n=2 replicates, since one of the replicates did not pass the QC (RIN <7) and was removed from the study.

## CONCLUSIONS

By studying how *M.tb* adapts to E-ALF, we were able to provide potential mechanisms explaining how E-ALF-exposed *M.tb* could escape into the cytosol by altering the expression of genes associated with many different virulence mechanisms such as ESX-secretion systems, Sec secretion systems, PDIMs metabolism, and ROS defense strategies, among others. Future functional studies will be required to delineate how these mechanisms potentially contribute to the susceptibility of the elderly population to TB pathogenesis.

## METHODS

### Human subjects and ethics statement

Human subject studies were conducted following stringent guidelines established by the US Code of Federal and Local Regulations (OSU Institutional review board IRB numbers 2012H0135 and 2008H0119). After receiving informed written consent, bronchoalveolar lavage fluid (BALF) from healthy adults (18 to 45 years old) and elderly (≥69 years old) participants from both sexes were collected. There was no consideration given to race or ethnicity. The following comorbidities were excluded: smokers (current or less than one month), excessive alcohol users, non-injection/injection recreational drug use, active pneumonia, asthma, bilateral cancer and on chemotherapy, Chronic obstructive pulmonary disease (COPD), pre-diabetes/diabetes (hemoglobin A1c higher than 5.7% within the last three months or a fasting blood glucose level of higher than 110 mg/dL), Body mass index (BMI) equal or higher than 40, hepatitis, human immunodeficiency HIV/AIDS, immunosuppression or taking nonsteroidal anti-inflammatory agents (TNF antagonists), leukemia/lymphoma, liver failure, renal failure, rheumatoid arthritis, pregnancy (or gave birth less than three months ago), taken antibiotics recently and/or regular treatment with over the counter anti-inflammatory medications.

### Human ALF isolation

Human BALF from healthy young and elderly donors was concentrated to obtain ALF at its physiological concentration within the human lung (1 mg/mL of phospholipid) as we reported in detail (10, 13–15, 24, 81).

### AT culture

The human ATs type II-like cell line A549 (ATCC^®^ CCL-185^TM^), a lung cancer cell line with many AT type II cell features (used as a model of ATs), was used for all cell culture experiments. Cell cultures were prepared as described (15, 24) with minor changes. Briefly, the A549 cell line was cultured at 37 °C with 5% CO_2_ in culture media (DMEM/F12 supplemented with 10% FBS [Atlas Biologicals, Fort Collins, CO]) and 1% PenStrep (Sigma, St. Louis, MO). One week before *M.tb* infection, cells were kept in antibiotic-free growth media.

### M.tb culture and exposure to human ALF

*M.tb* Erdman GFP strain (kindly provided by Dr. Marcus Horwitz, UCLA) was grown as described (10). Single cell bacterial suspensions of 1 × 10^9^ bacteria were generated as we reported (13, 15, 24) and exposed to a 50 μL pool of A-ALF (n=4 different human donors) and E-ALF (n=4 different human donors) for 12 h at 37°C with 5% CO_2_ (**Suppl. Fig. S1**). Following exposure, bacteria were thoroughly washed to remove any unbound ALF; the bacterial pellet was either stored at −80°C for further RNA extraction [after incubation in RNAProtect (Qiagen, Hilden, Germany) for 10 min at room temperature, RT] or suspended in culture media for AT infection. To assess *M.tb* viability and specifically to demonstrate that there are no differences in viable bacterial counts between E-ALF *vs*. A-ALF-exposed *M.tb*, ALF-exposed *M.tb* inoculums were serially diluted in 7H9 broth and plated on 7H11 agar supplemented with OADC (oleic acid-albumin-dextrose-catalase) enrichment. Unexposed *M.tb* were processed in parallel and used as control (bacterial pellet incubated for 12 h at 37°C without ALF). Three independent experiments were performed (prior to and during ATs infection); three different exposures to the same pools of ALF, using single bacterial suspensions of newly fresh harvested bacteria grown in 7H11 agar. Experimental conditions are shown in **Fig. 1A**.

### M.tb infection of Ats

*M.tb* infection of ATs was performed as we described in detail elsewhere (15, 24), using a multiplicity of infection (MOI) of 100:1 in 4 x 150 cm^2^ flasks per condition (**Suppl. Fig. S1**), which maintains an intact AT monolayer. After 2 h of infection, infected ATs were rinsed with culture media and cultured for up to 72 h post-infection (hpi) in culture media supplemented with 10 g/mL of gentamicin for subsequent downstream transcriptome analysis. Experimental conditions are shown in **Fig. 1B**. The chosen time point and MOI provide the quantity of bacterial RNA within ATs (as non-professional phagocytes) required to conduct the analysis described in this work. The stability of the infected monolayer was verified by microscopy before RNA extraction.

### M.tb RNA extraction, library prep, and RNA sequencing

After ALF exposure, bacterial pellets (before ATs infection) were incubated in RNAProtect (Qiagen, Hilden, Germany) for 10 min at RT, centrifuged, and stored at −80°C until further processing. RNA extraction was performed using the Quick-RNA Fungal/Bacterial Miniprep kit (Zymo Research, Irvine, CA, USA), following the manufacturer’s instructions and as we described (82). Briefly, bacterial pellets were suspended in RNA lysis buffer and transferred to a ZR BashingBead Lysis tube containing 0.1 mm and 0.5 mm ceramic beads. Samples underwent a bead beating procedure in a Disruptor Genie (10 cycles of 1 min at maximum speed with 1 min intervals on ice), facilitating the breaking down of the *M.tb* cell envelope. RNA in the supernatant was isolated using the Zymo-spin columns, including an in-column DNAse I treatment, and eluted in nuclease-free water. A second DNAse treatment was then performed using TURBO DNAse (Thermo Fisher Scientific, Waltham, MA, USA). RNA was further purified using the RNA Clean & Concentrator kit (Zymo Research, Irvine, CA, USA), and RNA concentration and quality were assessed using the Qubit 4 Fluorometer with the HS RNA kit (Thermo Fisher Scientific, Waltham, MA, USA). RIN numbers were acquired using a 4200 TapeStation System (Agilent, Santa Clara, CA, USA).

Infected ATs (72 hpi) were washed twice with culture media and then lysed with GTC solution (4M guanidinium thiocyanate, 0.5% sodium N-lauryl sarcosine, 25 mM tri-sodium citrate, 0.1M 2-mercaptoethanol, 0.5% Tween 80, pH 7.0) (69) for 5 min at RT and centrifuged at 5,000 x *g* for 30 min at RT to recover bacteria. *M.tb* pellets were washed in GTC lysis solution to combine the content from all flasks (4 x 150 cm^2^ flasks/ condition) into one single tube, centrifuged, and suspended in RNA lysis buffer. RNA was then extracted using the Quick-RNA Fungal/Bacterial Miniprep kit, treated with TURBO DNAse, and purified with the RNA Clean & Concentrator kit, as detailed above.

A whole-stranded transcriptome RNA library was generated using the Zymo-Seq RiboFree Total RNA kit (Zymo Research, Irvine, CA, USA), following the manufacturer’s protocol. RNA (250 ng) was used as input to construct the library, which included a rRNA depletion step of 30 min followed by 15 cycles of PCR. Using a NextSeq 500 Mid Output Illumina platform, a 75-bp paired-end (PE) sequencing was conducted (**Fig. 1C**). All samples were sequenced in the same flow cell to avoid sequencing batch effects. RNA sequencing was performed on three independent experiments (n = 3 per condition), except for E-ALF-exposed *M.tb* before infection (n=2), where one of the replicates was not included in the study due to low RNA concentration and quality (RIN < 7).

### Data analysis

After RNA sequencing, resulting reads were trimmed and aligned to the *M.tb* H_37_R_v_ reference genome (Genbank NC_000962.3) (83–85) with the STAR aligner v2.5.3a in the Partek Flow Genomic Analysis software (v9.0.20.0202, Partek Inc., Chesterfield, MO, USA), using default settings. Raw read count data were analyzed in iDEP.94 (http://bioinformatics.sdstate.edu/idep94/) (86). Briefly, data were filtered to remove poorly expressed genes. Differentially expressed genes (DEGs) that were significant between experimental conditions (**Fig. 1**) were identified using DESeq2 (87) with the following settings: Fold change (FC) of 2 (or log_2_ FC equal or greater than an absolute value of 1), and FDR cut-off of 0.1. The Benjamini-Hochberg procedure was used to adjust the false discovery rate (FDR), and fold changes were calculated using an Empirical Bayes shrinkage approach (87). Functional categories for DEGs were extracted from Mycobrowser (https://mycobrowser.epfl.ch/) (88). Venn diagrams were generated in Venny 2.1.0 (https://bioinfogp.cnb.csic.es/tools/venny/index.html) (89) or GraphPad Prism Version 9.5.1 Heatmaps were generated in GraphPad Prism Version 9.5.1.

## DECLARATIONS

### Ethics approval and consent to participate

Human subject studies were conducted following stringent guidelines established by the US Code of Federal and Local Regulations (OSU Institutional review board IRB numbers 2012H0135 and 2008H0119). Informed written consent was obtained before performing the bronchoalveolar lavage.

### Consent for publication

Not applicable.

### Availability of data and materials

All raw and processed sequencing data generated in this study was submitted to the NCBI Gene Expression Omnibus (GEO; https://www.ncbi.nlm.nih.gov/geo/) (90) under accession number GSE244851.

### Authors’ contributions

Study design and conceptualization, A.M.O.-F. and J.B.T.; Experimental procedures, A.M.O.-F., A.A.-G., and A.G.-V.; Acquisition of data and formal analyses, A.M.O.-F. and A.A.-G.; Collect BAL fluid, R.E.M and S.H.W; RNA library preparation and initial filtering/alignment of sequencing reads, J.G.; Writing manuscript, A.M.O.-F. and J.B.T.; Visualization and BioRender figures, A.M.O.-F; Writing—review and editing, A.M.O.-F., A.A.-G., J.B.T., J.T., Y.W. and L.S.S; Funding acquisition, A.M.O.-F., J.T., and J.B.T. All authors provided comments/edits and approved the final version of this manuscript.

## Acknowledgements

We thank Dr. Marcus Horwitz (University of California, Los Angeles) for kindly providing the GFP-*M.tb* Erdman strain. We acknowledge Clinton Christensen at the Texas Biomed Molecular Core and the BSL-3 Operations Program at Texas Biomed for their services and support.

