## Supplementary figures for "Impact of the elderly lung mucosa on *Mycobacterium tuberculosis* metabolic adaptation during infection of alveolar epithelial cells"

### FIGURE S1

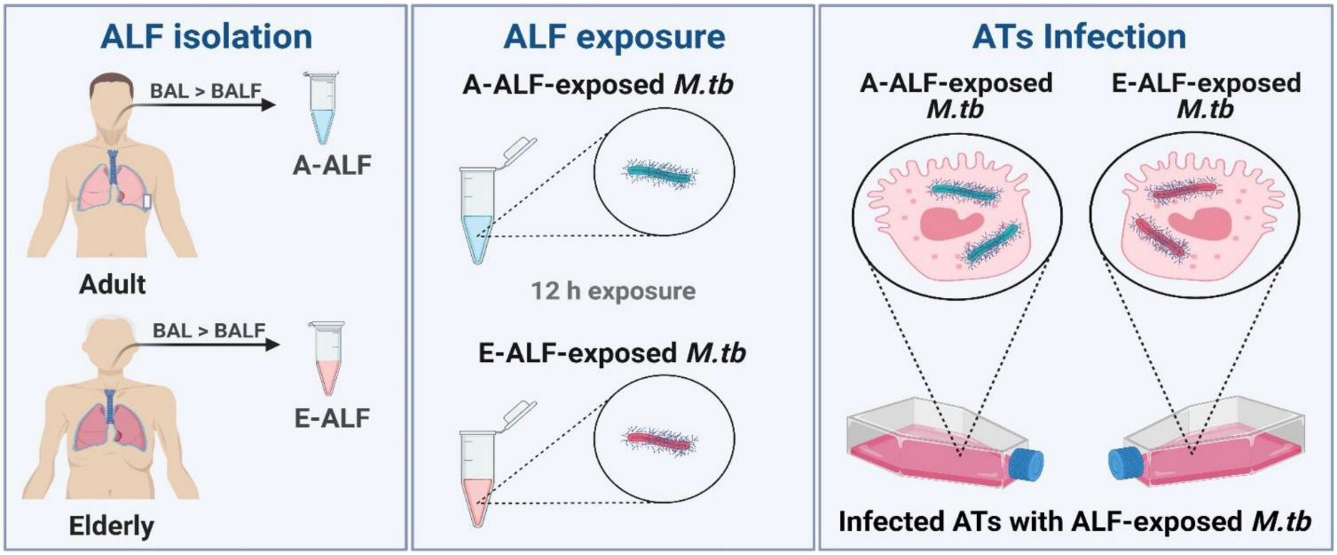


**Figure S1. Illustration of the experimental procedures performed in this study.** Isolation of ALF from Adult (A-ALF) and Elderly (E-ALF) individuals, followed by A- or E-ALF exposure of *M.tb* and subsequent infection of ATs. Figure created using BioRender (https://biorender.com/).

**FIGURE S2**

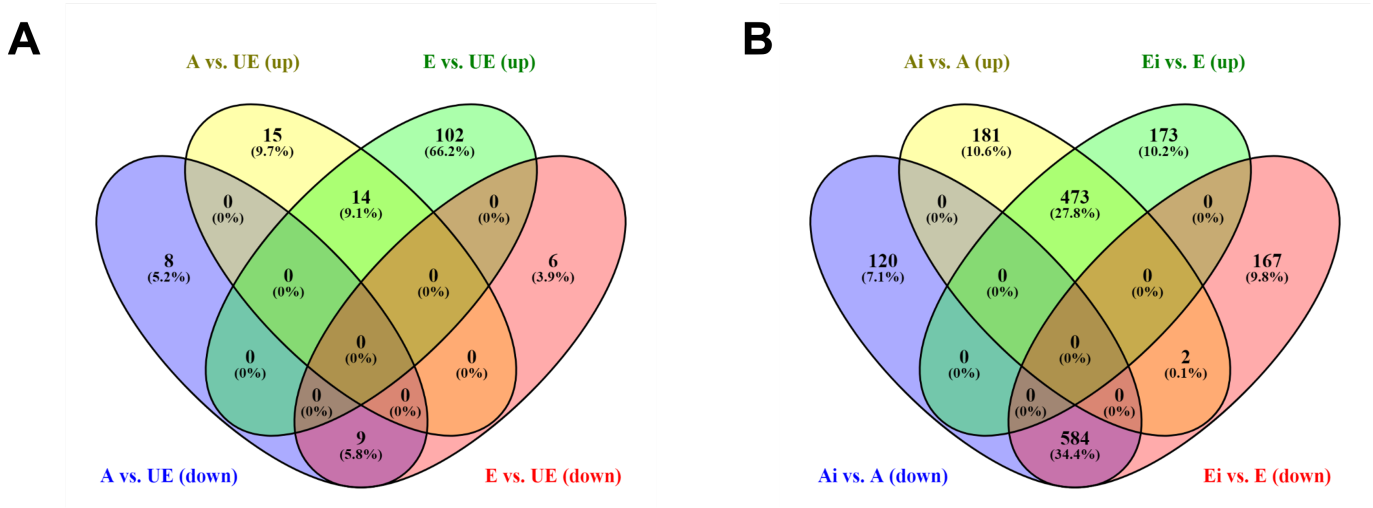


**Figure S2. Venn diagram of DEGs of ALF-exposed *M.tb*.** Unique and shared DEGs of ALF-exposed *M.tb* prior to (**A**) and during ATs infection (**B**). Venn diagrams were generated using Venny 2.1.0. A: A-ALF-exposed *M.tb*; E: E-ALF-exposed *M.tb*; UE: unexposed *M.tb*; Ai: A-ALF- exposed *M.tb* during ATs infection; Ei: E-ALF-exposed *M.tb* during ATs infection.

**FIGURE S3**

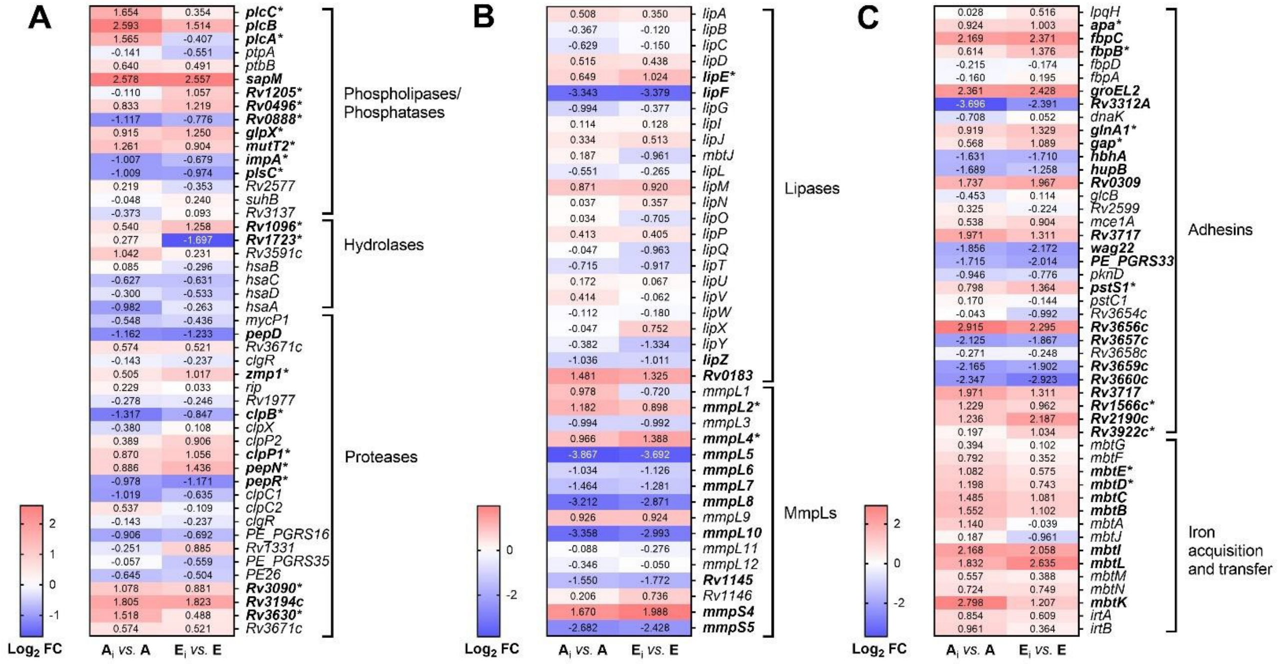


**Figure S3. Heatmaps of *M.tb* genes associated with virulence factors at 72 hpi in ATs.** (**A**) Phospholipases/Phosphatases, Hydrolases and Proteases; (**B**) Lipases and MmpL; and (**C**) Adhesins during *M.tb* replication in ATs. Cells depict Log2 FC values in ALF-exposed *M.tb* in ATs ($A_i$: Adult ALF-exposed *M.tb* during infection; $E_i$: Elderly ALF-exposed *M.tb* during infection) *vs.* ALF- exposed *M.tb* before infection (A: Adult ALF; E: Elderly ALF), upregulated: red, downregulated: blue. Genes in bold indicate significant DEGs (Log2 FC equal or greater than an absolute value of 1, FDR < 0.1) for both conditions. Genes with an asterisk indicate a significance in only one of the comparisons, highlighting differences between conditions. Note the different scales used in A, B, and C for better visualization of the results . Heatmaps were generated in GraphPad Prism v9.1.1.

### SUPPLEMENTARY TABLES

**Table S1. Complete list of *M.tb* DEGs (upregulated and downregulated) after exposure to ALF before and during ATs infection at 72 hpi**. DE is shown as Log2 FC; up- and downregulated genes in orange and blue, respectively; significant DEGs (Log2 FC equal or greater than an absolute value of 1 and FDR < 0.1) are highlighted in bold. DEGs are listed based on locus tag, and a description of functional category is included (Mycobrowser and Uniprot).

**Table S2. DE analysis of *M.tb* genes (all genes) after exposure to ALF before and during ATs infection at 72 hpi.** DE is shown as Log2 FC; significant DEGs (Log2 FC equal or greater than an absolute value of 1 and FDR < 0.1) are highlighted in bold. DEGs are listed based on locus tag, and a description of functional category is included (Mycobrowser and Uniprot).
